# Genomic architecture of migration timing in a long-distance migratory songbird

**DOI:** 10.1101/2022.09.14.508039

**Authors:** Evelien de Greef, Alexander Suh, Matt J. Thorstensen, Kira E. Delmore, Kevin C. Fraser

**Affiliations:** Department of Biological Sciences, University of Manitoba, Winnipeg, Canada R3T 2N2; Department of Organismal Biology, Uppsala University, Uppsala, Sweden SE-752 36; School of Biological Sciences, University of East Anglia, Norwich, United Kingdom NR4 7TU; Department of Biology, Texas A&M University, College Station, Texas 77843

## Abstract

The impact of climate change on spring phenology poses risks to migratory birds, as migration timing is controlled predominantly by endogenous mechanisms. Despite numerous studies on internal cues controlling migration, the underlying genetic basis of migration timing remains largely unknown. We investigated the genetic architecture of migration timing in a long-distance migratory songbird (purple martin, *Progne subis subis*) by integrating genomic data with an extensive dataset of direct migratory tracks. Our findings show migration has a predictable genetic basis in martins and maps to a region on chromosome 1. This region contains genes that could facilitate nocturnal flights and act as epigenetic modifiers. Additionally, we found that genomic variance explained a higher proportion of historic than recent environmental spring phenology data, which may suggest a reduction in the adaptive potential of migratory behavior in contemporary populations. Overall, these results advance our understanding of the genomic underpinnings of migration timing and could provide context for conservation action.

## Introduction

Climate change affects spring phenology in temperate zones and could have significant, negative impacts on migratory animals [1]. For example, migrants must synchronize arrival at the breeding grounds to coincide with seasonal resources [2]. These resources are becoming available earlier and it is unclear if migrants will be able to match these advances, potentially leading to substantial population declines [3]. Migratory timing is largely endogenously controlled [4], and thus knowledge of its genetic architecture (e.g., the identity, number, and location of genetic loci involved) is essential for predicting if and how migrants will respond to phenological changes that accompany climate change.

Previous genetic studies provide important insights regarding migration timing, such as in genes associated with circadian and circannual rhythms [5,6]; however, results vary across species [7] and are limited to small portions of the genome. Another limitation associated with earlier studies was an inability to quantify migratory behavior in the wild—prior to 2007, it was not possible to track animals <100 g on migration [8] and, for example, most migratory avian species fall into this size class. We overcame these limitations here, combining high resolution genomic data with an extensive migration tracking dataset for purple martins (*P.s. subis*). The purple martin is a Nearctic-neotropical migrant that travels over 7,000 km between North America and South America [9] and exhibits extensive latitudinal variation in migration timing. It is thus a powerful system to study migration genomics. For example, individuals breeding in the southern edge of the range in Florida may arrive as early as mid-January, while their northern counterparts in Alberta may arrive as late as June [10].

Our first objective was to examine the genomic architecture of migration timing by assembling a reference genome for the purple martin and integrating sequencing data with light-level geolocator tracks. We examined results from genome-wide association studies (GWAS), polygenic scores (PGS), and genomic differentiation analyses. Our second objective was to examine the adaptive potential of migration timing by comparing genomic variation associated with historic and contemporary spring phenology. We reran our GWAS using environmental proxies for spring timing and tested if the proportion of phenotypic variation explained by our genomic data (hereafter “PVE”) was lower in contemporary datasets, which could suggest a change in adaptive potential. This study expands our understanding of the whole-genome contribution to migration and yields insight into the adaptability of migration timing behavior.

## Results

### Reference and resequencing data

The final P. *subis* reference genome assembly based on long reads and linked reads was 1.17 Gb in length, consisted of 2,896 scaffolds, had an N50 scaffold length of 6.13 Mb and an N50 contig length of 3.08 Mb. The annotation included 12,686 genes (SI Appendix, Table S1). The assembly length was similar to other avian genomes, which are typically between 1.0–1.2 Gb [11]. BUSCO analysis revealed that the P. *subis* genome was relatively complete with 91% of avian orthologs detected as complete sequences (89.1% being single-copy and 1.9% being duplicated), which was in range of other non-model avian genomes [12]. We aligned resequencing data for 87 individuals to this reference resulting in 4.6 million SNPs after filtering. All these individuals were tracked on migration with light-level geolocators yielding precise estimates for migratory timing.

### Genomic architecture of migration timing

Birds in this study exhibited considerable latitudinal variation in migratory timing (sampling locations in Table S2), ranging over 120 to 131 days for spring departure and arrival dates (Figure S1). Spring departure and arrival locations are displayed in Figure 1, showing weak migratory connectivity between breeding and wintering sites (i.e. mixing of breeding populations at shared wintering areas) such as observed in Fraser et al. 2012 and Fraser et al. 2017 [9,13]. Estimates of PVE from Bayesian sparse linear mixed models (BSLMMs) [14] were high, with a median value of 0.70 (89% ETI: 0.09–1.00). PGS estimated from linear mixed models [15] were strongly correlated with spring timing (*R*^2^ of 0.25, *p* = 1.4e^-148^). Through jackknife cross-validation partitions, we assessed predictive power of the PGS model and found that birds with lower PGS deciles exhibited earlier migration timing compared with individuals in higher PGS deciles with later migration timing (Figure 2a). BSLMMs did not identify any specific genomic regions linked to migratory timing, except for one SNP with PIP = 0.18 (Figure S2), 79 kb upstream of gene *tsc-22*. However, a survey of net genomic differentiation (ΔF_ST_) between the earliest and latest spring migrants in our dataset did reveal a region of elevated differentiation on chromosome 1 (Figure 3). ΔF_ST_ controls for processes unrelated to migration that could elevate F_ST_ (including population structure, see Methods). This elevation was additionally present in comparisons of early and late migrants within populations in Florida (southernmost colony) and Alberta (northernmost colony) (Figure 3b) suggesting population structure did not generate this pattern. Reductions in nucleotide diversity and Tajima’s D indicative of a selective sweep are also present in this region, which covers 2 Mb region and consist of 13 genes (Table S3) including *ppfia2* and *nts*, which may be related to sleep [16,17], and *mettl25* and *acss3* that may serve as important epigenetic modifiers [18,19].

**Figure 1.**
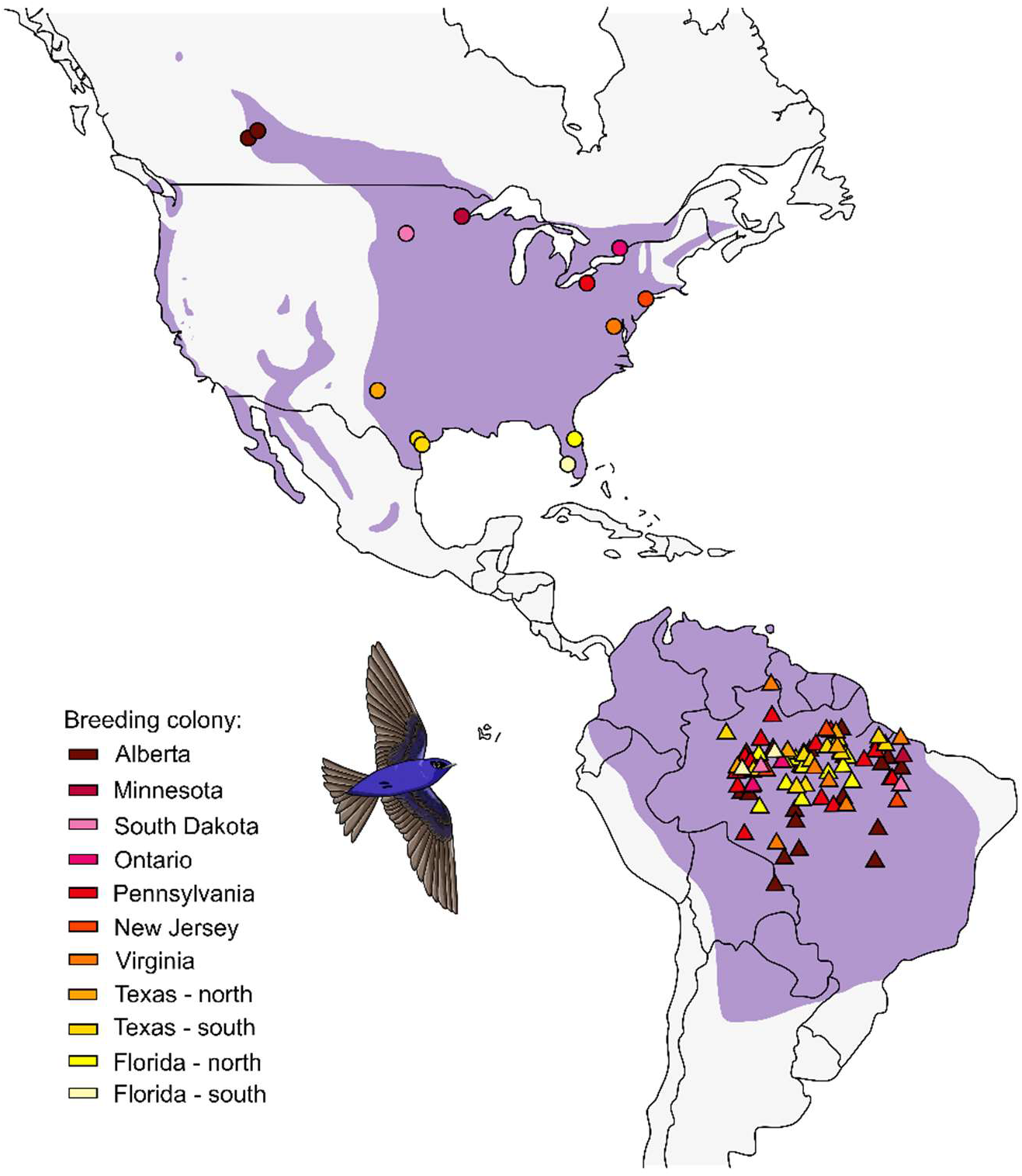
Purple martin breeding and wintering distribution (purple), including sampling sites for 87 individuals in their North American breeding range (circles) and their respective South American wintering destination before spring departure (triangles). Individuals from distinct breeding colonies overlapping at the wintering grounds demonstrate weak connectivity between breeding and wintering sites (such as observed in Fraser et al. 2012, 2017).

**Figure 2.**
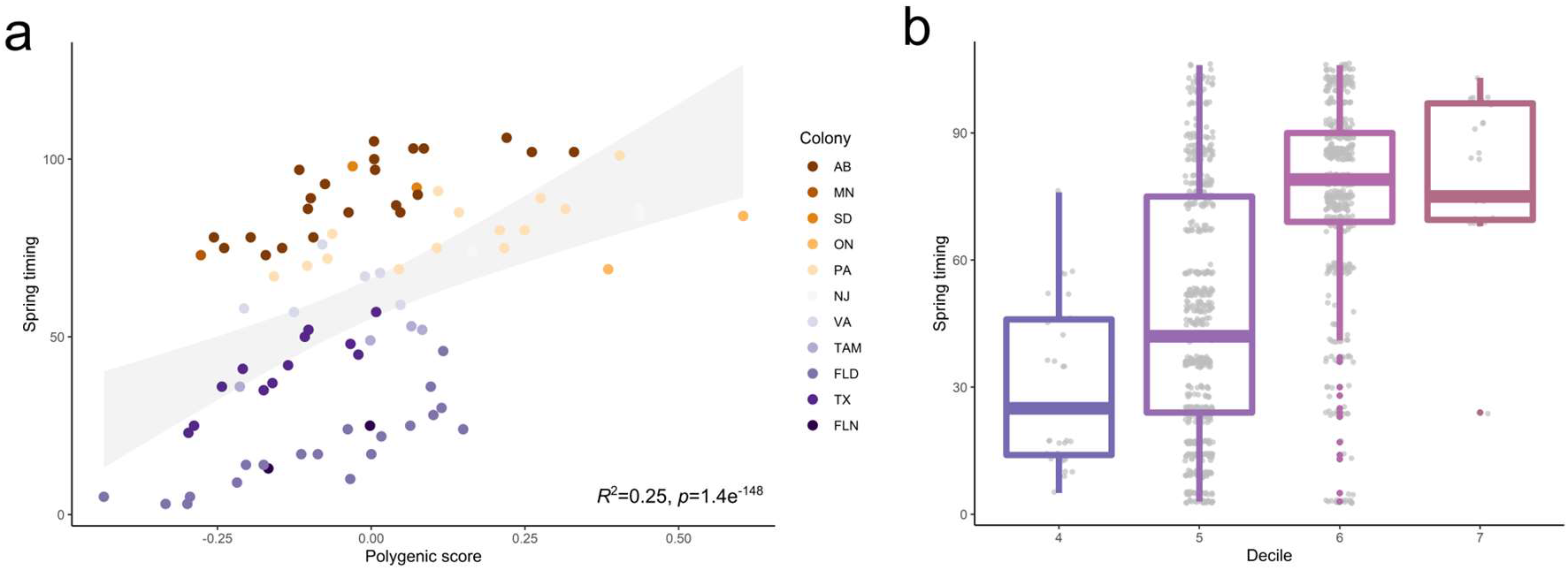
(a) Polygenic scores of spring migration timing for purple martins (n = 87) colored in order by latitude, and linear regression standard error is colored in gray. (b) Individuals in lowest decile of predicated polygenic scores (PGS) had earlier migration timing compared with individuals in higher deciles with later timing.

**Figure 3.**
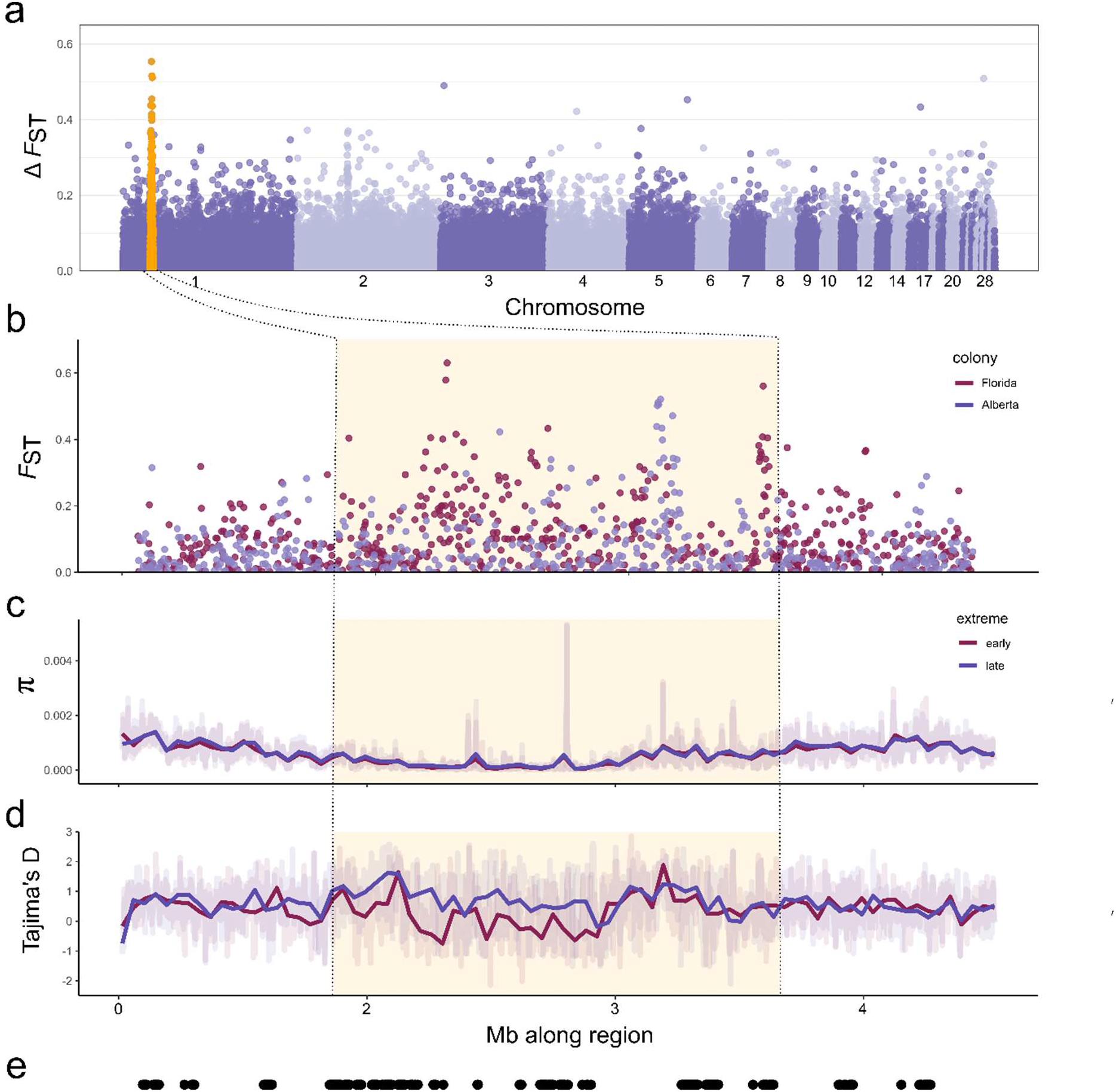
(a) Net genetic differentiation (ΔF_ST_) across autosomes in 5 kb non-overlapping windows between earliest and latest spring migrants. The elevated region on chromosome 1 is highlighted in orange, with plots examining this region to show (b) F_ST_ within Alberta and within Florida, (c) nucleotide diversity (π), (d) Tajima’s D, and (e) location of genes in this region as black dots.

### Genomic association with ecological spring indices

We extracted green-up [20] and first bloom [21] data for sites where purple martins were tracked. Green-up data were available for all sites (US & Canada; 2001–2015); first bloom data were available for US sites only but spanned a longer period (1981–2015). There was a strong correlation between these environmental variables and spring migration timing (*R* = 0.76–0.90), thus we used these variables as proxies for current and historic migration timing in our analyses. Estimates of PVE were consistently higher in the historic datasets (green-up PVE = 0.98, SD = 0.06, 89% ETI = 0.91–1.00; first-bloom PVE = 0.99, SD = 0.07, 89% ETI = 0.91–1.00), compared to PVE in recent years (green-up PVE = 0.79, SD = 0.3, 89% ETI = 0.07–1.00; first bloom PVE = 0.89, SD = 0.16, 89% ETI = 0.52–1.00) (Figure S3).

## Discussion

We used one of the largest tracking datasets available for a long-distance migratory songbird and a genome-wide SNP dataset to reveal the genetic underpinnings of migration timing. Our results demonstrate a strong genetic basis to migration timing in the purple martin. We discovered a previously unidentified 2 Mb genetically differentiated region on chromosome 1, illuminating components underlying migration timing. Additionally, lower PVE with recent environmental data suggest a reduced adaptive potential with advancing spring phenology.

Phenotypes that vary across individuals are a result of both environmental and genetic factors, and PVE represents the proportion of variance attributed to genetic factors. The large PVE estimate (0.7) is evidence that variation in migration timing is largely determined by genetics. The predictive utility of the polygenic model across multiple deciles showed it is possible to predict early and late migrants using genetic variants, which could potentially aid in estimating a birds’ phenotype in the wild. With overlapping wintering grounds among purple martin colonies [9], the model may also help predict an individuals’ timing tendency in addition to breeding region when captured during the winter. While the high genomic variation explaining migratory timing does not preclude phenotypic plasticity, it suggests that changes in timing may occur through microevolutionary processes. It is important to understand the source of considerable genetic variation in migratory traits [24], and future work will inform how influences such as standing genetic variation (presence of more than one allele at a locus in a population) may play a role in facilitating rapid microevolutionary changes [25]. While our sample size limits our estimates of PVE (the ETIs were quite wide), our polygenic scores (PGS) supported prediction of individual migration timing. Given the present sample size, it is remarkable that a substantial proportion of variation was explained through genomics. Whether this proportion could be even greater with larger sample sizes [22,23] could be determined in future studies. However, collecting enough samples to capture strong power in tests of genomic associations with phenotypes in field-based wildlife research will be challenging.

The small number of genomic loci in the GWAS significantly associated with spring migration suggest this trait is controlled by many alleles of small effect, which may have been undetected, or could have been located in assembly gaps [11]. However, elevated genomic differentiation on chromosome 1 illuminates a potential connection between migration timing in purple martins with some genes related to rest and epigenetic modifiers. *Ppfia2* and *nts* are 2 of the 13 genes located in this region. *Ppfia2* has been linked to sleep and wakefulness in white-crowned sparrows [16], and *nts* has been linked to sleep regulation in European mice [17]. While purple martins are primarily diurnal migrants, they can incorporate both day and night flights on spring migration [26], and the former genes could play a role in these nocturnal flights. *Mettl25* and *acss3* could mediate epigenetic changes in response to environmental cues important for migratory timing; *mettl25* encodes a methyltransferase that represses gene expression [18,19] and *acss3* produces acetyl-CoA which promotes gene expression by acetylating histones. Acetyl-CoA is also important for generating, using, and storing energy [27]. While this study suggests associations with these genes, further work could elucidate these mechanisms and their roles in migration timing. While estimates of F_ST_ are often considered bottom-up comparisons, we compared extreme phenotypes while controlling for population differentiation to identify genomic regions associated with migration timing. Results from this approach were further supported by a comparison of early and late migrants within populations that recovered a similar elevated pattern of genomic differentiation.

Our comparison of contemporary and historical data on spring phenology suggests that the adaptive potential of migration timing in purple martins may have declined in recent years, with lower values of PVE in contemporary datasets. These reductions could derive from selection for earlier arrival on the breeding grounds and will ultimately affect the amount of genetic variation available for future change. Interpretation of these results requires caution (e.g., our analyses assume phenotypic plasticity has not changed, we are using environmental variables as a proxy for migration timing and are using data from a small subset of populations). In addition, environmental and genetic differences between the present day and when historical spring phenology data were recorded (1981–1984 and 2001–2004) may have affected the interactions between genotype, phenotype, and environment in ways not captured by the samples collected more recently (2008 – 2015). Therefore, generalizability of GWAS and PVE over time cannot be rigorously assessed with the present data. Nevertheless, the presence of this pattern in our data set could be a signal of a broader underlying change in adaptive potential which should be further investigated.

This study presents novel findings on migration timing, opening the door to understanding components of the genomic architecture of migration timing in other long-distance migrants. The strong genomic variation and significant regions associated with purple martin migration timing could have important implications for adaptability in long-distance migrants. If the genomic potential for adaptability has decreased in recent years, this could hinder the ability of migrants to keep up with the pace of changing climates. Many portions of the genome are conserved across other bird species and vertebrates [28] and climate change continues to affect migratory animals all over the world. Therefore, these findings bring us closer to understanding a common basis for migration, which may have broad implications for a variety of organisms.

## Methods

### Reference and resequencing data

The reference genome was assembled using PacBio long and 10X linked reads generated for a female martin from Manitoba, Canada. We used FALCON [29,30] to create the initial assembly, then polished and scaffolded the genome with ArrowGrid, Pilon, and ARKS [31,32,33]. Then we annotated the genome through MAKER [34]. We used skimSeq (low-coverage whole-genome sequencing) to generate resequencing data [35] for an additional 87 birds (average coverage 2.7x per sample). Missing genotypes were then imputed with Beagle [36], using information from the reference, surrounding genotypes, linkage disequilibrium structure, and haplotype blocks [37]. These included 45 male and 42 female blood samples collected from 13 different breeding colonies across North America between 2008–2015 (Table S2). We filtered SNPs for quality (QUAL>20, MQ>20), max-missing (20%), minor allele frequency (MAF>0.05), Hard-Weinberg equilibrium, and biallelic sites. Details on assembly, annotation, sequencing, and filtering are in supplementary information.

### Light-level geolocator analysis

Light-level geolocators were mounted during the breeding season using leg-loop backpack harnesses and retrieved through recapture in the following year. Purple martin behavior of aerial foraging and use of open habitats makes light-level geolocators ideal for capturing sunrises and sunsets with minimal shading. The timing of these twilights is used to estimate the daily locations of birds over the entire year, using the midpoint of rise-set events to determine longitudes and day length for estimating latitudes [38]. We analyzed twilight times with BAStag and GeoLight [39,40], producing estimated daily locations to obtain migratory departure and arrival dates. Due to the correlation of departure with arrival timing for migratory journeys, we ranked individuals in order of timing for both dates and combined these values to determine overall timing phenotypes for spring migration.

### Genomic architecture of migration timing

BSLMMs and LMMs were run using GEMMA [14], where we included the covariates of sex, year, age, and the first principal component (PC1) from a PCA summarizing genetic variation in our dataset. We summarized results from these runs for PVE and used posterior inclusion probabilities (PIP) to identify specific SNPs with strong associations to the timing phenotypes. PIP is the probability that the SNP is associated with the phenotypic variation [41] and following [42] we considered SNPs with PIPs > 0.1 important. Polygenic models were created using the PLINK v1.9 [43] and following Choi et al. (2020)’s PGS pipeline [44]. We used VCFtools [45] to estimate F_ST_ between the 10 earliest (originating from two Florida colonies) and 10 latest (originating from two Alberta colonies and one Virginia colony) spring migrants. Since this F_ST_ could be elevated by processes unrelated to migration, including linked background selection and population structure [46], we controlled for these potential effects by subtracting F_ST_ between Alberta and Florida (representing the northernmost and southernmost breeding regions) from values estimated between extreme timing phenotypes. This approached has been used in crows [47] and blackcaps [48] to isolate differentiation associated with specific phenotypes. Additionally, we estimated F_ST_ between early and late migrants within Alberta and Florida populations separately to examine if we could recover the same signature of elevated F_ST_ in the same genomic region.

### Genomic association with ecological spring indices

Green-up data were extracted from MODIS [20] and first bloom dates from the USA National Phenology Network [21]. We extracted data for each purple martin colony location over all available years. We ran BSLMMs for “historic” (2001-2004 for green-up and 1981-1984 for first bloom) and recent phenology data (up till 2015) and ran BSLMMs for each association test, using year, PC1, and colony as covariates. PC1 controls for population structure and colony accounts for the fact that birds from the same colony are assigned the same values for each environmental variable each year. The green-up (MODIS) association used all 87 purple martin individuals, and the first bloom (NPN) dates spanned 63 purple martin individuals (USA only).

## Acknowledgements

We thank K. Applegate, J. Barrow, L. Burgess, L. Chambers, T. Cheskey, P. Clifton, B. Dietrich, J. Fischer, G. Hvenegaard, P. Kramer, P. Mammenga, N. Mickle, M. North, M. Pearman, J. Ray, A. Ritchie, A. Savage, T. Shaheen, J. Siegrist, C. Silverio, B. Stutchbury, and J. Tautin for sample collection; M. Przeworski, Z. Fuller, and C.J. Garroway for conceptual input; F. Taborsak-Lines, M. Ormestad, P. Ewels, C. Wang, O.V. Pettersson, M.B. Mosbech, and C. Tellgren-Roth for generating data for the reference assembly; K. Jeffries, National Genomics Infrastructure (NGI)/Uppsala Genome Center, and Texas A&M AgriLife Genomics and Bioinformatics Service for laboratory support; Texas A&M High Performance Research Computing and Compute Canada for computing services. This work was supported by funding from the University of Manitoba, Texas A&M University, Natural Sciences and Engineering Research Council, Research Manitoba, the Society of Canadian Ornithologists, Sigma Xi, RFI/VR & Science for Life Laboratory, Sweden, and we further acknowledge the Knut and Alice Wallenberg Foundation and the Swedish Research Council, and SNIC/Uppsala Multidisciplinary Center for Advanced Computational Science for assistance with massively parallel sequencing and access to the UPPMAX computational infrastructure.

## Author contributions

K.F. and K.D. designed and supervised the study. K.F. conducted/coordinated the collection of migration data and blood sampling. E.D. conducted laboratory work and performed genomic analyses with K.D.’s guidance. M.T. assisted with association modelling and conducted polygenic score analyses. A.S. coordinated sequencing for the reference genome and provided guidance on the assembly. E.D. analyzed light-level geolocator data K.F.’s guidance. E.D., K.F., and K.D. wrote the manuscript with feedback from M.T. and A.S.

## Data availability statement

Raw sequencing data is available on SRA NCBI BioProject PRJNA772931. Source code for reference genome and resequencing work is available at https://github.com/edegreef/PUMA-reference-genome and https://github.com/edegreef/PUMA-resequencing-data.

