## supplementary information for "Genomic architecture of migration timing in a long-distance migratory songbird"

### Methods

#### Reference genome

*Genome assembly*

Data for the reference genome came from a second-year female purple martin from Manitoba, Canada (49°44' N, 97°7' W). DNA from this individual's blood was extracted using an in-situ agarose plug extraction procedure at the Science for Life Laboratory in Sweden. Pacific Biosciences (PacBio) long-read libraries [1] and 10X Genomics Chromium (10X) linked-read libraries were prepared from this DNA [2], and sequenced on a PacBio Sequel instrument (5 SMRT cells) and an Illumina HiSeq X instrument (1 lane, 2x 150 bp paired-end reads), respectively. The initial reference was assembled with the PacBio sequences using FALCON algorithms with SMRT Link v7.0.0 [3,4]. We polished the genome using two rounds of the program ArrowGrid v0.6.0 with the PacBio subreads bam files [5]. To prepare for the next polishing step, we aligned 10X fastq files to the arrow-corrected genome using BWA v0.7.17 [6] and sorted the resulting bam file with SAMtools v1.9 [7]. We then polished the genome with Pilon v1.23 using 10X reads to correct bases, fixed mis-assemblies, and filled gaps [8]. We scaffolded the genome with ARKS v1.0.4 [9], using interleaved 10X reads assembled with LongRanger v2.2.2 [10]. We checked the genome for duplicated scaffolds using dedupe.sh in BBMap v38.44 [11]. To look for possible contaminants, we used the nucleotide database from the National Center for Biotechnology Information [12] and the program BLAST+ v2.9 [13] to identify any foreign DNA. The final genome was evaluated with QUAST v5.0.2 [14], Assemblathon 2 [15], and BUSCO v4.0.5 with the aves dataset (aves_odb10) and augustus species set to chicken [16].

*Genome annotation*

We annotated the genome using four rounds of the program MAKER v2.31.10 [17]. The first round included RepeatMasker and Exonerate using protein data from three model species: chicken (*Gallus gallus*; assembly GRCg6a), collared flycatcher (*Ficedula albicollis*; assembly FicAlb_1.4), zebra finch (*Taeniopygia guttata*; assembly bTaeGut1_v1) obtained from Ensembl database [18]. A hidden Markov model (HMM) was created using the outputs from the first round. This HMM was then used in the second round to train SNAP using computational gene prediction. The third round used the updated HMM model from the second round and included the Augustus chicken model. The fourth and final round had the same parameters as the third, but additionally filtered out genes of annotation edit distance (AED) score over 0.5. After completing the MAKER rounds, the annotation was aligned with protein data from the uniprot database [19] using BLAST+ v2.9 [13] and protein functions from InterProScan v5.4 [20].

Resequencing data

*Sample collection and DNA extractions*

All birds were captured with drop door traps while feeding their young at their nesting colonies in artificial housing. Sampling locations and numbers are listed in Table S2. These samples were collected between 2008–2015. Blood collection and DNA extraction followed the same methods seen in de Greef et al. 2022 [21], where up to 150 µl of blood was drawn from each bird's brachial vein and stored in Queen's Lysis buffer and DNA was extracted using the Qiagen DNeasy Blood & Tissue kits.

*Sequencing & Imputation*

We used a low-coverage whole-genome approach (skimSeq) to generate resequencing data for these birds [22-24], following the methods of de Greef et al. 2022 [21]. Specifically, libraries were prepared with NEXTFLEX Rapid XP DNA-SEQ Kit and sequenced on Novaseq 6000 platform (2x 150bp paired-end reads) to 3x coverage. This approach for assaying the entire genome with low coverage has been validated in the 3K rice genome project which simulated low coverage sequencing from high quality high coverage public data [25,26]. Our data were filtered and trimmed using Trimmomatic v0.38 [27] then aligned to the reference genome using Bowtie2 v2.3.4.2 [28]. These alignments were sorted, realigned, and filtered (MQ>5) using SAMtools v1.9 [7] and PICARD v.2.18.4 [29]. The variants were called in GATK v3.5 using HaplotypeCaller [30]. Missing genotypes were imputed with Beagle v.4.0 [25], using information from surrounding genotypes, linkage disequilibrium structure, and haplotype blocks [22]. imputation performance was evaluated by masking a random 3% of genotypes as missing then imputed, which resulted in 0.9 accuracy with a 0.9 genotype probability cut-off. SNPs were filtered with this 0.9 genotype probability threshold.

*SNP filtering*

We filtered the final set of SNPS using vcflib v1 [31] and VCFtools v.0.1.16 [32], limiting analyses to SNPs with quality score (QUAL) > 20 and mapping quality (MQ) > 20. We removed sites with indels, and then filtered the SNPs to a dataset with minor allele frequency (MAF) > 0.05 (5%), max-missing genotype score of 0.2, matching Hardy-Weinberg equilibrium, and containing only biallelic sites. We removed scaffolds with abnormally high read depths (4 or higher), as they likely represent repetitive regions. We also removed Z and W-linked scaffolds to generate an autosomal dataset. These sex-linked scaffolds were identified from mapping the genome to a chicken genome with SatsumaSynteny [33] and by using $F_{ST}$ values through VCFtools v.0.1.16 [31] between males and female samples. We used a PCA on to control for population structure in our GWAS. We ran the PCA using smartpca in the Eigensoft program [34], using SNPs that were pruned for linkage disequilibrium with PLINK v1.9 [35].

Genome-wide association studies

*Light-level geolocator analysis*

To define sunrises and sunsets, we used the preprocessLight function in the R-package BAStag [36]. Here, the light-intensity threshold was set at 32 to systematically separate day and night, and false twilights or outliers due to shading or light pollution were removed through manual inspection. In addition, the sudden onset of extreme light fluctuations in spring indicated the bird entering and exiting nest cavities, signifying the spring arrival date. Next, we estimated daily coordinates using the twilight data with the coord function in R-package GeoLight [37]. Here, the calibration for the sun elevation angle was calculated using two weeks of light data at the end of the breeding season, where the bird was known to stay in their respective colony before departing on fall migration. Since there is little variation in day lengths around the spring and fall equinox, the latitudes around the equinox were removed using a tol of 0.13 [37] as these would result in inaccurate locations. We determined spring departure and arrival dates from the location coordinates based on sudden shifts in latitude, and confirmed with the changeLight function, which defined residency periods throughout the year.

*Association Mapping*

BSLMMs were run in GEMMA [38] after converting the SNP dataset from a vcf file to a binary plink format (bim, bed, fam files) using PLINK v1.9 [35]. GEMMA includes uses a kinship matrix to control for factors that influence phenotypes and are correlated with genotypes (e.g., population structure). We also included several additional covariates in our model to control for these effects: sex, year, age, and the first principal component (PC1) from a PCA summarizing genetic variation in our dataset. PC1 was the only significant component in our PCA and showed only weak clustering by latitude with 1.5% variance explained. BSLMMs do not permit the inclusion of covariates and thus we regressed these covariates out of the timing phenotypes and used the residuals as the corrected phenotype for our models. We ran four independent chains for each BSLMM, with a burn-in of 5 million steps and a subsequent 20 million MCMC steps. In the spring BSLMM we found one SNP with PIP = 0.18 near gene *tsc-22*.

*Polygenic score*

LMM results obtained in GEMMA [41] were used to estimate polygenic scores (PGS) using covariates sex, year, age, PC1, colony, distance travelled, and duration. Polygenic models were created using the PLINK v1.9 [35] and following Choi *et al.* (2020)'s PGS pipeline [42]. To assess the predictive power of the PGS model, we used jackknife cross-validation partitions and randomly sampled from the overall dataset with a training/test dataset of an 85/15 split for 100 iterations [43]. We examined the accuracy in terms of $R^2$ in the training data and plotted PGS deciles.

*Differentiation across genome*

Mixed models run in GEMMA did not identify any specific genomic regions linked to migration timing. This result could suggest there is a polygenic basis to migratory timing or our sample size was too small. Accordingly, we took a second approach to identify genomic variants underlying timing, estimating differentiation ($F_{ST}$) across the genome between the earliest and latest migrants ($n$ = 10 for each extreme phenotype). Earliest migrants included samples from Lacombe, Alberta (n = 6), Camrose, Alberta (n = 2), Virginia (n = 1), and Pennsylvania (n = 1). Latest migrant included samples from Naples, Florida (n = 1) and Bay Lake, Florida (n = 9). $F_{ST}$ could be elevated by processes unrelated to migration, including linked background selection and population structure [44]. We controlled for these potential effects by subtracting $F_{ST}$ between Alberta and Florida (representing the northernmost and southernmost breeding regions) from values estimated between extreme timing phenotypes. This approach has been used in crows [45] and blackcaps [46] to isolate differentiation associated with specific phenotypes. Additionally, $F_{ST}$ estimated between early and late migrants within Alberta (n = 8) and Florida (n = 8) separately was also elevated in this region, further supporting that this region is related to migration timing as opposed to population structure. We used VCFtools v.0.1.16 [32] to estimate $F_{ST}$, nucleotide diversity, and Tajima's D. Pairwise $F_{ST}$ across sampling groups in the study ranged between regions ranged from 0.009 (Alberta & Florida) to 0.001 (Texas & Florida)

<u>Genomic association with ecological spring indices</u>

Green-up data, representing the onset of greenness, was extracted from MODIS using the MCD12Q2 Land Cover Dynamics V005database [47]. We obtained first bloom through the USA National Phenology Network, consisting of the average first bloom dates for Red Rothomagensis lilac (*Syringa x chinensis*), Arnold Red honeysuckle (*Lonicera tatarica*), and Zabelii honeysuckle (*Lonicera korolkowii*) [48]. To categorize "historic" and recent phenology data, we grouped the oldest years (2001–2004 for green-up, 1981–1984 for first bloom) and recent years with a four-year interval to the year the blood was sampled, and kept the interval between each group consistent. We ran BSLMMs as described above, including year, PC1 and colony as covariates. PC1 controls for population structure and colony accounts for the fact that birds from the same colony are assigned the same values for each environmental variable each year. There was no significant difference between time frames in the variance exhibited by these variables indicating that results from spring phenology analyses are unlikely to be driven by change in variances between time periods.

**Table S1**. Purple martin (*P.s. subis*) draft reference genome assembly metrics

| **Assembly metrics** | |
|---|---|
| Genome size (bp) | 1,165,951,862 |
| Number of scaffolds | 2,896 |
| Longest scaffold (bp) | 45,082,031 |
| Shortest scaffold (bp) | 16,249 |
| Mean scaffold length (bp) | 402,608 |
| N50 scaffold length (bp) | 6,129,949 |
| N50 contig length (bp) | 3,084,734 |
| L50 scaffold count | 44 |
| GC content (%) | 43.06 |
| N's per 100 kb | 3.99 |
| Complete BUSCOs | 7592 (91.0%) |
| Complete single-copy BUSCOs | 7431 (89.1%) |
| Complete duplicated BUSCOs | 161 (1.9%) |
| Fragmented BUSCOs | 118 (1.4%) |
| Missing BUSCOs | 628 (7.6%) |
| Number of identified genes | 12,686 |

**Table S2**. Sampling locations for 87 geolocator-tracked purple martins.

| Province/State | Colony | Latitude | Longitude | N | Year(s) |
|---|---|---|---|---|---|
| Alberta | Camrose | 53.011 | -112.864 | 5 | 2012-2014 |
| Alberta | Lacombe | 52.391 | -113.612 | 17 | 2012-2014 |
| Florida | Bay Lake | 28.360 | -81.588 | 19 | 2013-2015 |
| Florida | Naples | 26.149 | -81.746 | 2 | 2013 |
| Minnesota | Millie Lacs | 46.146 | -93.724 | 1 | 2011 |
| New Jersey | Middletown | 40.390 | -74.001 | 3 | 2011 |
| Ontario | Ottawa | 45.351 | -75.827 | 2 | 2014 |
| Pennsylvania | Erie | 42.115 | -80.145 | 14 | 2008-2013 |
| South Dakota | Columbia | 45.598 | -98.310 | 2 | 2011-2013 |
| Texas | Amarillo | 35.040 | -101.933 | 4 | 2013 |
| Texas | Corpus Christi | 27.680 | -97.407 | 4 | 2013 |
| Texas | Sandia | 27.997 | -97.879 | 8 | 2013 |
| Virginia | Woodbridge | 38.613 | -77.263 | 6 | 2010-2013 |

**Table S3.** List of genes in elevated $F_{ST}$ region on chromosome one.

| scaffold | position start | position end | gene | description |
|---|---|---|---|---|
| scaffold110 | 46198 | 72361 | SLC6A15 | Solute Carrier Family 6 Member 15 |
| scaffold110 | 660079 | 668953 | CEPT1 | Choline/ethanolamine phosphotransferase 1 |
| scaffold110 | 1317685 | 1336461 | METTL25 | Methyltransferase like 25 |
| scaffold110 | 1336917 | 1356721 | CCDC59 | Coiled-coil domain containing 59 |
| scaffold110 | 1767838 | 1859397 | PPFIA2 | PTPRF interacting protein alpha 2 |
| scaffold110 | 1899350 | 1967674 | ACSS3 | Acyl-CoA synthetase short chain family member 3 |
| scaffold22 | 74157 | 151868 | ALX1 | ALX homeobox 1 |
| scaffold22 | 205263 | 213135 | CEPT1 | Choline/ethanolamine phosphotransferase 1 |
| scaffold22 | 366758 | 368017 | RASSF9 | Ras association domain family member 9 |
| scaffold22 | 419805 | 431374 | NTS | Neurotensin/neuromedin N |
| scaffold22 | 464152 | 471608 | MGAT4C | MGAT4 family member C |
| scaffold22 | 1228676 | 1250364 | C12orf50 | Chromosome 12 open reading frame 50 |
| scaffold22 | 1252756 | 1257492 | C12orf29 | Chromosome 12 open reading frame 29 |

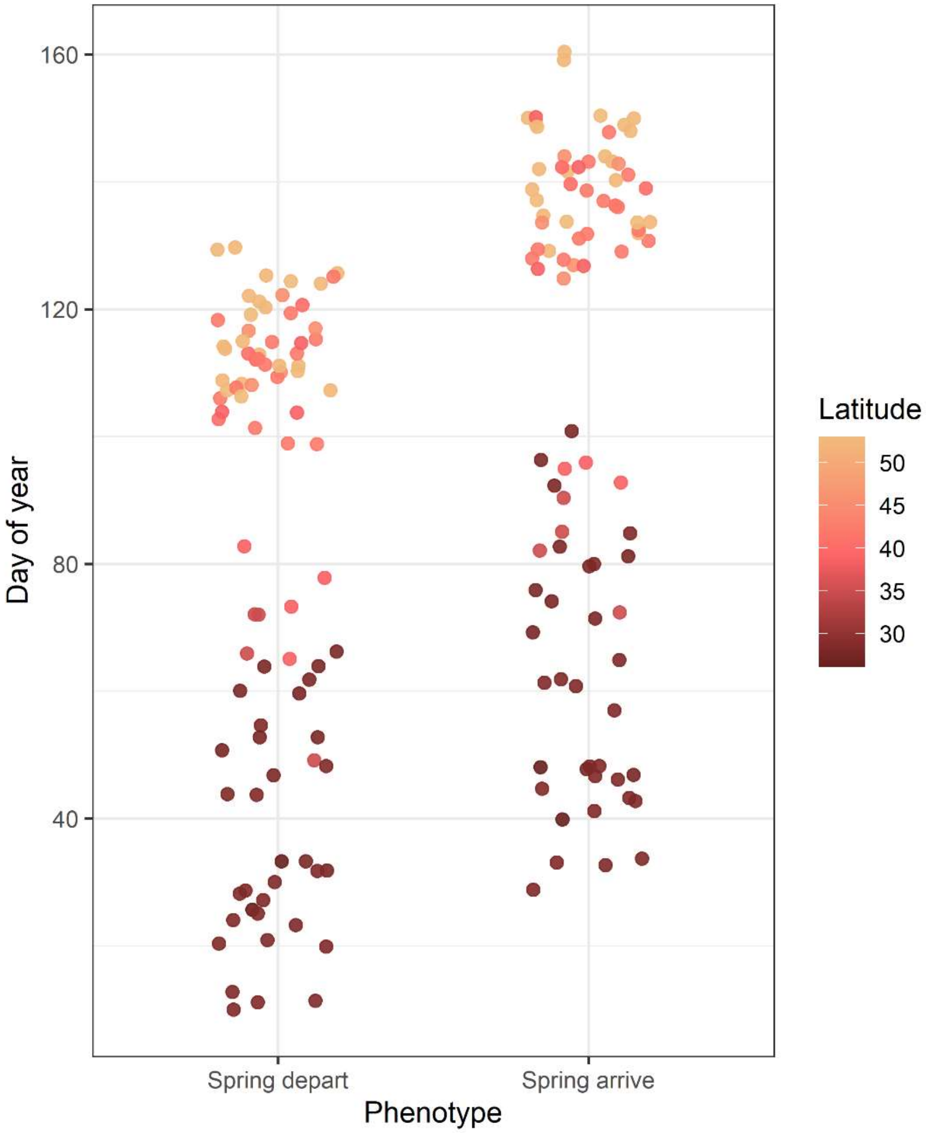


**Figure S1**. Migratory timing phenotypes of 87 geolocator-tracked purple martins, including the start and end of spring migration in Julian date format for each bird. Migration timing follows a latitudinal trend, where individuals breeding at higher latitudes exhibit later timing.

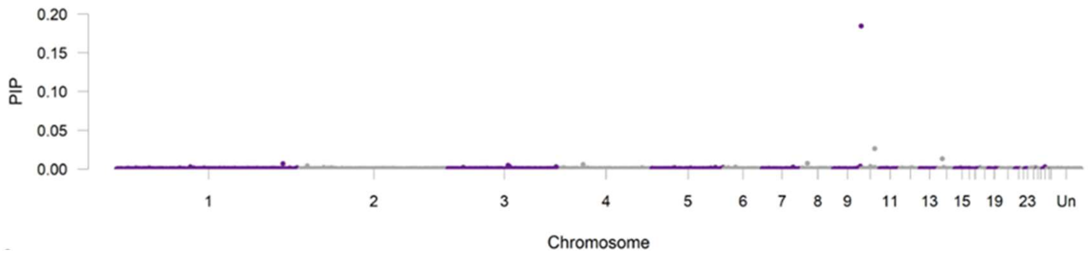


**Figure S2**. Bayesian sparse linear mixed model of spring migration

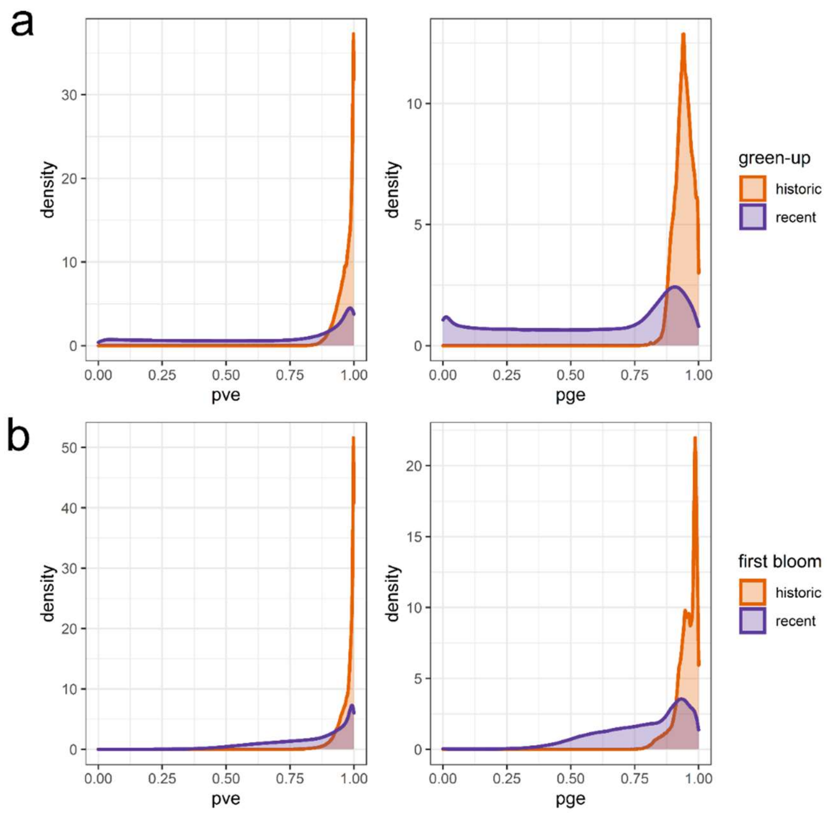


**Figure S3.** PVE and PGE estimates comparing “historic” and recent years with a) green-up data between 2001-2011, and b) first bloom data between 1981-2015. Both display overall higher PVE and PGE in older time periods and lower PVE and PGE in more recent years.
